# DNA-intercalating antiphage molecules trigger abortive infection through ‘mutual destruction’ and synergize with bacterial immunity

**DOI:** 10.64898/2026.01.13.699316

**Authors:** Larissa Ernst, Cornelia Gätgens, Bente Rackow, Nadiia Pozhydaieva, Elyès Gaaloul, Aileen Krüger, Johannes Seiffarth, Michelle Bund, Vivien Joisten-Rosenthal, Dietrich Kohlheyer, Björn Usadel, Alexander Harms, Katharina Höfer, Julia Frunzke

## Abstract

Bacteria deploy diverse antiphage defense systems, including small bioactive molecules providing protection at the multicellular level. DNA-intercalating anthracyclines, such as daunorubicin, exhibit broad antiphage activity, but the underlying mechanism has remained elusive. Here, we systematically screened the *Escherichia coli* BASEL phage collection to elucidate the mode of action of DNA-intercalating antiphage molecules. We identified taxonomically distinct clusters of susceptible viral groups and show that, in T5-like phages (*Markadamsvirinae*), daunorubicin blocks infection after first-step transfer (FST). In the presence of daunorubicin, continued expression of pre-early genes leads to abortive infection via ‘mutual destruction’, where both phage and host succumb. Analogous abortive-infection phenotypes occur across taxonomically diverse phages exposed to chemically distinct DNA-intercalating molecules. Notably, we show that daunorubicin synergizes with downstream nucleic acid-targeting defenses underscoring context-dependent outcomes. Together, these findings reveal how chemical defense contributes to the multilayered antiviral immunity and highlight the intricate interplay between mechanistic inhibition and infection outcome.

## Introduction

The ongoing evolutionary arms race between bacteria and phages drives a dynamic process of co-evolution. Bacteria have evolved a wide array of antiphage defense systems to counter viral predation, while phages, in turn, develop counter-measures to overcome these protective barriers (Bernheim & Sorek, 2018; Chevallereau et al., 2022; Koonin et al., 2006; Koskella & Brockhurst, 2014). The investigation of bacterial antiphage defense systems has emerged as a rapidly expanding research field, not only for advancing our fundamental understanding of virus-host interactions but also for the potential biotechnological exploitation of these systems as novel molecular tools. To date, an ever-growing number of defense mechanisms have been identified, extending beyond the classic nucleic acid-targeting restriction-modification (RM) systems and CRISPR-Cas to encompass a striking diversity of strategies, including abortive infection (Abi), nucleotide depletion, and cyclic-oligonucleotide-based anti-phage signalling systems (CBASS) (Georjon & Bernheim, 2023).

Abi had initially been introduced as a broad term describing the phenomenological category of antiphage defense systems inducing programmed cell death (PCD) after the detection of a viral infection (Lopatina et al., 2020). In recent years, numerous novel antiphage systems have been discovered that appear to function as a ‘last line of defense’ by triggering host cell death. Yet, emerging evidence suggests a more complex relationship between the underlying mechanism of defense and the resulting infection phenotype. In a recent opinion article, Aframian and Eldar argue for disentangling mechanism from phenotype. They explicitly distinguish between cell death via the classical PCD targeting host components and death of the host driven by the action of phage components even after successful inhibition of viral infection, a process they define as ‘mutual destruction’ (Aframian & Eldar, 2023).

In addition to RNA- and protein-based defense systems, small, bioactive molecules naturally produced by *Streptomyces* spp. have been shown to act as potent antiphage agents. Reported examples include in particular DNA-intercalating molecules, such as anthracyclines, as well as diverse molecules belonging to the class of aminoglycosides (Brock & Wooley, 1963; Kever et al., 2022; Kronheim et al., 2018; Morita et al., 1979; Parisi & Soller, 1964). Due to their secretion into the environment, this so-called chemical defense via small molecules bears the potential to confer protection against viral infection at the community level (Hardy et al., 2023; Kever et al., 2024).

Daunorubicin (Dau), also referred to as daunomycin, was previously reported to exhibit broad antiphage activity (Kronheim et al., 2018). This anthracycline is a well-established chemotherapeutic agent, primarily used to treat myeloblastic and acute lymphoblastic leukaemias. Its cytotoxic activity stems from its ability to inhibit human topoisomerase II through the intercalation of its planar group between adjacent DNA/RNA base pairs, thereby preventing the progression of nucleic acid replication (Brossard & Corcelli, 2024). DNA intercalating agents, which perturb DNA metabolism in both eukaryotic and prokaryotic cells (Brossard & Corcelli, 2024; Hind et al., 2019), represent promising candidates in the search for novel antiphage compounds. While their antiphage activity was recognized already in the past century (Morita et al., 1979; Parisi & Soller, 1964), it was only recently that they have been appreciated as part of the natural *Streptomyces* antiviral immune system. The mechanism by which daunorubicin inhibits phage infection and the factors determining viral susceptibility to daunorubicin - even at concentrations without significant effects on the host - have remained elusive. Previous work has shown that daunorubicin acts at an early stage of the phage replication cycle after DNA injection but prior to the onset of genome replication (Kronheim et al., 2018).

To unravel the mode of action of daunorubicin and to identify phage determinants conferring sensitivity to DNA-intercalating antiphage molecules, we harnessed the *Escherichia coli* BASEL (**Ba**cteriophage **Se**lection for your **L**aboratory) collection (Maffei et al., 2021). Systematic screenings revealed taxonomically-related clusters of phage families showing high susceptibility to the antiphage action of daunorubicin. Exemplified for T5-like phages (*Demericviridae* / *Markadamsvirinae*), we further demonstrate that antiphage activity is mediated through ‘mutual destruction’. Daunorubicin caused the blockage of phage infection after first-step transfer at the level of pre-early genes expression, which ultimately imposes a lethal outcome on the bacterial host. We further demonstrate that the defense phenotype is context dependent, as daunorubicin synergizes with downstream nucleic acid–targeting systems. These results emphasize the intricate interaction between different lines of defense ultimately defining bacterial antiviral immunity.

## Results

### Taxonomically distinct pattern of daunorubicin sensitivity among phage families

Previous studies have identified the DNA-intercalating compound daunorubicin as a broad-spectrum antiphage agent that blocks an early step of infection (Kronheim et al., 2018). To elucidate its mode of action and uncover phage-encoded determinants of sensitivity, we leveraged the *Escherichia coli* BASEL phage collection, which enables systematic analyses across a wide taxonomic diversity of phages (Maffei et al., 2021). As bacterial host, we utilized the *E. coli* K-12 MG1655 ΔRM strain lacking all native restriction-modification systems (ΔRM), the abortive infection systems PifA and RexAB as well as the O-antigen glycan barrier. This strain provides a genetically simplified background, minimizing confounding interactions with native bacterial immunity and allowing for a more direct assessment of daunorubicin’s antiphage activity (Maffei et al., 2021).

Exposure of the phage collection to increasing concentrations (0-20 µM) of daunorubicin during infection of *E. coli* K-12 MG1655 ΔRM revealed distinct sensitivity patterns (Figure 1A). Notably, members of the *Drexlerviridae* phage family exhibited markedly reduced efficiency of plating (EOP) at 10 µM daunorubicin. Similar sensitivity was observed for members of the *Demerecviridae* family and the *Vequintavirinae* subfamily. In contrast, an intermediate level of susceptibility was noted for the *Seuratvirus* genus within the *Queuovirinae* subfamily. Conversely, phages belonging to the *Dhillonvirus* genus, the entire *Tevenvirinae* subfamily, and the *Autographviridae* family displayed pronounced resistance to daunorubicin in this host genetic background. Notably, neither GC content nor phage genome size showed significant correlation with the observed daunorubicin sensitivity patterns (Figure 1B,C).

**Figure 1:**
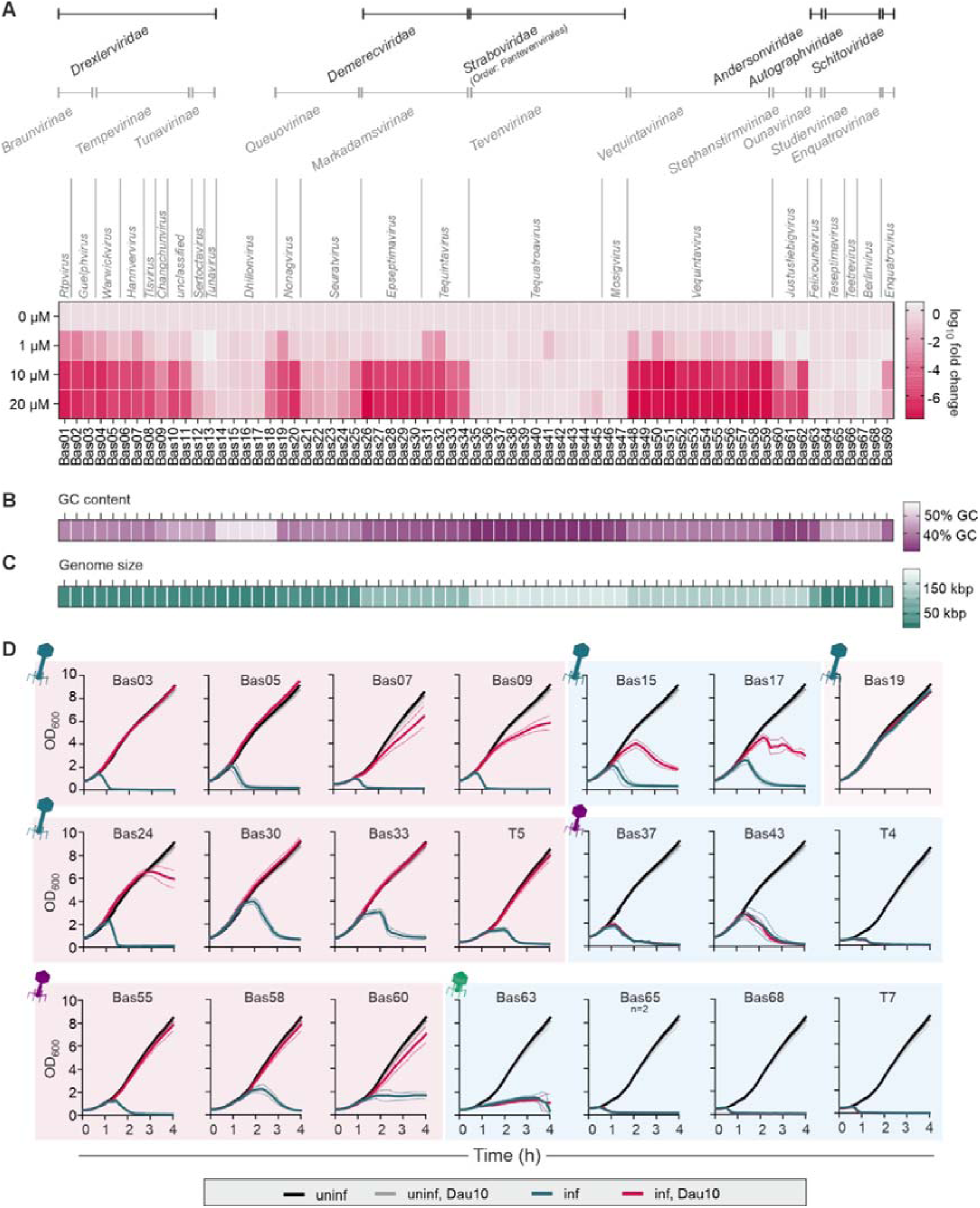
Screening of the BASEL phage collection revealed taxonomically distinct patterns of phages with high sensitivity to daunorubicin. A) Heat map showing the log_10_ FC in efficiency of plating (EOP) upon addition of increasing daunorubicin concentrations to LB double-agar overlay assays inoculated with in E. coli K-12 MG1655 ΔRM (n = 2). B) GC-content and C) genome size of the different BASEL phages (Maffei et al. (2021). D) Liquid infection experiments of selected phages as representatives of the different taxonomic groups. Infection was performed at an MOI of 0.1 in LB medium with and without 10 µM daunorubicin (n = 3). Morphotypes of the different phage families are indicated with respective icons (blue: siphovirus, purple: myovirus, green: podovirus). Phages showing sensitivities towards daunorubicin in the plate screening are shaded red, whereas resistant phages are shaded blue.

These findings were further validated using infection assays in liquid cultures at a multiplicity of infection (MOI) of 0.1 in the presence of 10 µM daunorubicin (Figure 1D). Except for Bas19, which caused no growth defect upon phage addition even in absence of daunorubicin, infection dynamics were largely consistent with the results obtained on plate screenings. Phages classified as daunorubicin-sensitive caused reduced or even no detectable host cell lysis under drug exposure, while phages from the resistant *Tevenvirinae* and *Autographviridae* displayed no significant alteration in infection efficiency. Interestingly, although *Dhillonvirus* phages Bas15 and Bas17 were classified as resistant, they exhibited a mild yet reproducible reduction in host lysis in liquid infection assays, indicating partial susceptibility to daunorubicin without complete inhibition of phage propagation.

### High resistance of Tevenvirinae phages is not conferred through genome hypermodification

Apart from genome size and GC content, covalent base modifications could be another factor influencing the antiviral activity of DNA intercalating compounds. The daunorubicin-resistant *Tevenvirinae*, including the prominent *E. coli* model phage T4, are known for their large genome size of >160 kbp and their high resistance to DNA-targeting immune mechanisms such as restriction-modification systems. This resistance is attributed to the hypermodification of cytosines by hydroxymethyl glycosylation for the *Tequatrovirus* genus and hydroxy arabinosylation for the *Mosigvirus* genus (Mahler et al., 2025; Thomas et al., 2018; Weigele & Raleigh, 2016). To test whether these base modifications affect infectivity in presence of daunorubicin, we compared the infection dynamics of T4 and T4 Δα-/β-gt (Δgt), which lacks the α-/β-glycosyltransferases responsible for the final glycosylation of hydroxylmethyl-dCTPs (provided by Marianne De Paepe). In presence of 20 µM daunorubicin, no difference in infectivity was observed between the tested conditions for either T4 variant (Figure S1B,C) on *E. coli* K-12 MG1655 ΔRM. Based on these results, we concluded that the DNA modification does not severely impact the phage susceptibility to daunorubicin.

### Daunorubicin causes an abortive infection phenotype

Daunorubicin efficiently inhibits infection of complete families of phages such as *Drexlerviridae* or *Demerecviridae* (Figure 1). To gain insights into the defense phenotype, infection was performed at low and high MOIs of 0.25 and 2.5, respectively (Figure 2A). In the case of the sensitive phages Bas07 and Bas09 of the *Braunvirinae* subfamily (of *Drexlerviridae* family), as well as Bas33 and T5 of the *Markadamsvirinae* subfamily (of *Demerecviridae* family), infection at a low MOI of 0.25 resulted in a delayed or even absent culture collapse after addition of daunorubicin (light red vs light blue line). Interestingly, infection in the presence of daunorubicin at an MOI of 2.5 transferred cells into a growth-arrested state (pink vs. dark blue line), which is indicative for an abortive infection phenotype. In the absence of daunorubicin, infection led to rapid culture clearance. In contrast, resistant phages such as T4 from the *Tevenvirinae* subfamily (family *Straboviridae*) and T7 from the *Autographviridae* family exhibited an identical infection dynamics, regardless of the presence of daunorubicin at either MOI.

**Figure 2:**
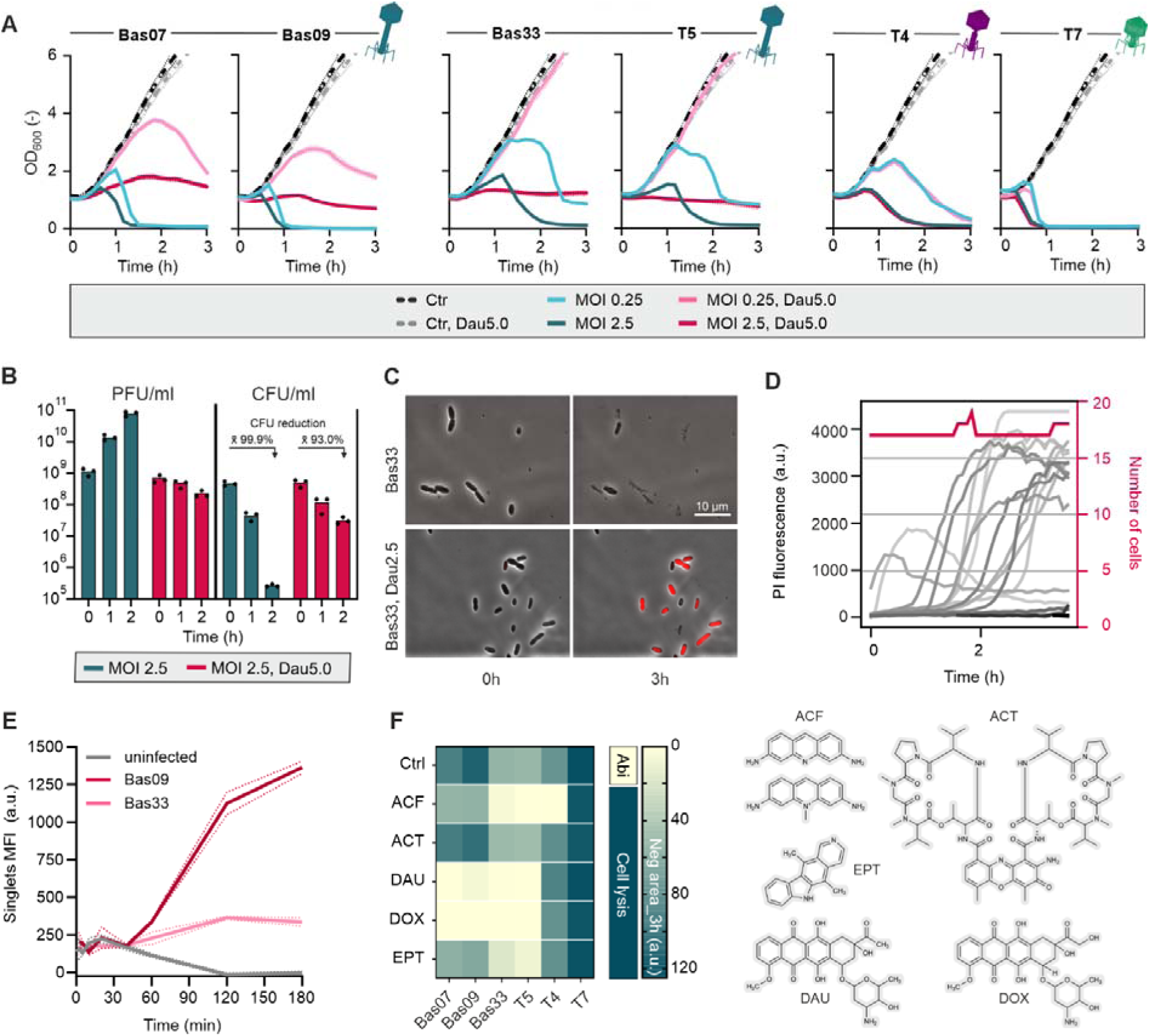
Infection in presence of daunorubicin causes an abortive infection phenotype. A) Liquid infection assays of selected phages in E. coli K-12 MG1655 ΔRM in presence and absence of 5 µM daunorubicin at low and high MOIs of 0.25 and 2.5, respectively. B) PFU and CFU counts upon infection with Bas33 +/- 5 µM daunorubicin. The mean values (x̄) of the percentage reduction in CFU/ml are shown. CFU counts were determined by spotting a decimal dilution series on LB agar plates containing 0.1% citrate for phage inactivation. C) Single-cell analysis of E. coli K-12 MG1655ΔRM infected with Bas33 (Vequintavirinae) in microfluidic chips +/- 5 µM daunorubicin upon addition of Basel phage Bas33. Complete time lapse videos are provided in Video S1 and S2. Propidium iodide staining was used to visualize increased membrane permeability and cell death. The same LUT settings were applied to all images. D) PI fluorescence and number of cells upon infection of E. coli K-12 MG1655 ΔRM with Bas33 in presence of 2.5 µM daunorubicin in microfluidic chips, shown for one representative single replicate. E) Uptake of daunorubicin by E. coli K-12 MG1655 ΔRM upon phage infection at an MOI of 2.5 in presence of 5 µM daunorubicin, measured as cell fluorescence via flow cytometry (488 nm laser). F) Heat map showing the negative area (a.u.) upon infection with selected Basel phages in presence of different DNA intercalating compounds (ACF: acriflavine, 5 µg/ml; ACT: actinomycin D; 5 µM; DAU: daunorubicin, 5 µM; DOX: doxorubicin, 5 µM; EPT: elipticine, 5 µM). Negative areas of ≤20 indicate an abortive infection phenotype, while increasing negative areas >20 indicate phage-induced cell lysis as for the compound-free control (ctrl) (n = 3). Structures of the respective intercalators are indicated.

Growth patterns aligned with plaque- (PFU) and colony-forming units (CFU) counts. While CFU declined with rising PFUs under normal infection conditions due to cell lysis and progeny release, no phage amplification occurred with daunorubicin, though CFUs still dropped by ∼93% after 120 min (Figure 2B). Time-resolved single-cell microscopy confirmed the growth stagnation, showing further an increased membrane permeability and subsequent cell death in the presence of daunorubicin, as indicated by the applied propidium iodide staining and the increased daunorubicin uptake at later stages of infection (Figure 2C, D, E, Figure S2).

In addition to analyzing the effect of daunorubicin on infection dynamics, we also examined the effect of other DNA-intercalating agents, including doxorubicin (anthracycline), acriflavine (acridine derivative), actinomycin D (polypeptide antibiotic), and ellipticine (tetracyclic alkaloid). Negative areas, defined as deviations of the growth curves below the baseline during a 180 min infection period, were quantified and classified as ≤ 20 (a.u.), indicative for an Abi phenotype, or > 20 (a.u.), indicative for cell lysis (Figure 2F, Figure S2). As expected, doxorubicin showed almost identical effects on phage infection as daunorubicin. Interestingly, ellipticine and acriflavine exhibited no phage-inhibiting activity against the *Drexlerviridae*, but against the *Demerecviridae* phages Bas33 and T5. Notably, despite resistance to all other tested compounds, infection of phage T4 was inhibited by acriflavine. Contrary to this, actinomycin D showed no activity against all tested phages and T7 showed no susceptibility to all tested intercalators.

### Daunorubicin blocks infection of T5-like phages after FST causing ‘mutual destruction’ via expression of pre-early genes

To gain deeper mechanistic insights, we determined transcriptomic and proteomic changes in the early stages of bacteriophage infection with and without daunorubicin treatment. We focused on the daunorubicin-sensitive T5-like phage Bas33, known to infect via a two-step delivery of the phage genome (Davison, 2015). RNA-Sequencing revealed distinct differences in the transcription profile of phage genes after 20 minutes of infection in the presence and absence of daunorubicin. While late genes coding for structural proteins, such as *bas33_0007* (major capsid protein) or *bas33_0014* (tail tube protein), were already actively transcribed under normal infection conditions, the presence of daunorubicin revealed a sharply defined expression pattern, with transcription largely restricted to the pre-early phage genes (Figure 3 A,B). These genes are located in the first 8% of the phage genome, which is injected during the first-step transfer (FST). Of these 17 pre-early genes, A1 of phage T5 was shown to be essential for host DNA degradation, while both A1 and A2 are essential for the second-step transfer (SST). The deoxyribonucleotide 5L monophosphatase (*dmp*) facilitates infection by enabling further degradation of host nucleotides during host takeover and preventing their incorporation into viral DNA (Davison, 2015). Zooming in into the log_2_ FC (Dau_5_/Dau_0_; FDR ≤ 0.05) of the pre-early phage region confirmed the comparable gene expression between both conditions, showing log_2_ FC below the threshold of ± 2 (Figure 3C). Time-resolved analysis of pre-early gene transcription via RT-qPCR revealed further that addition of daunorubicin led to a up to 4.5-fold reduction in transcripts of *A1 (bas33_0182)* and *A2 (bas33_0180)* within the first 10 min of transcription, while respective transcripts levels exceeded the ones of the control conditions at 20 min post infection. In contrast, early gene expression *(bas33_0147)* was only detected after 20 min, with transcripts levels being 20-fold higher in absence of daunorubicin (Figure 3D). Transcripts of the *E. coli* housekeeping gene *atpD* remained stable over the 20-minute time course, indicating that Bas33 does not rapidly degrade the host genome upon infection, in contrast to canonical T5.

**Figure 3:**
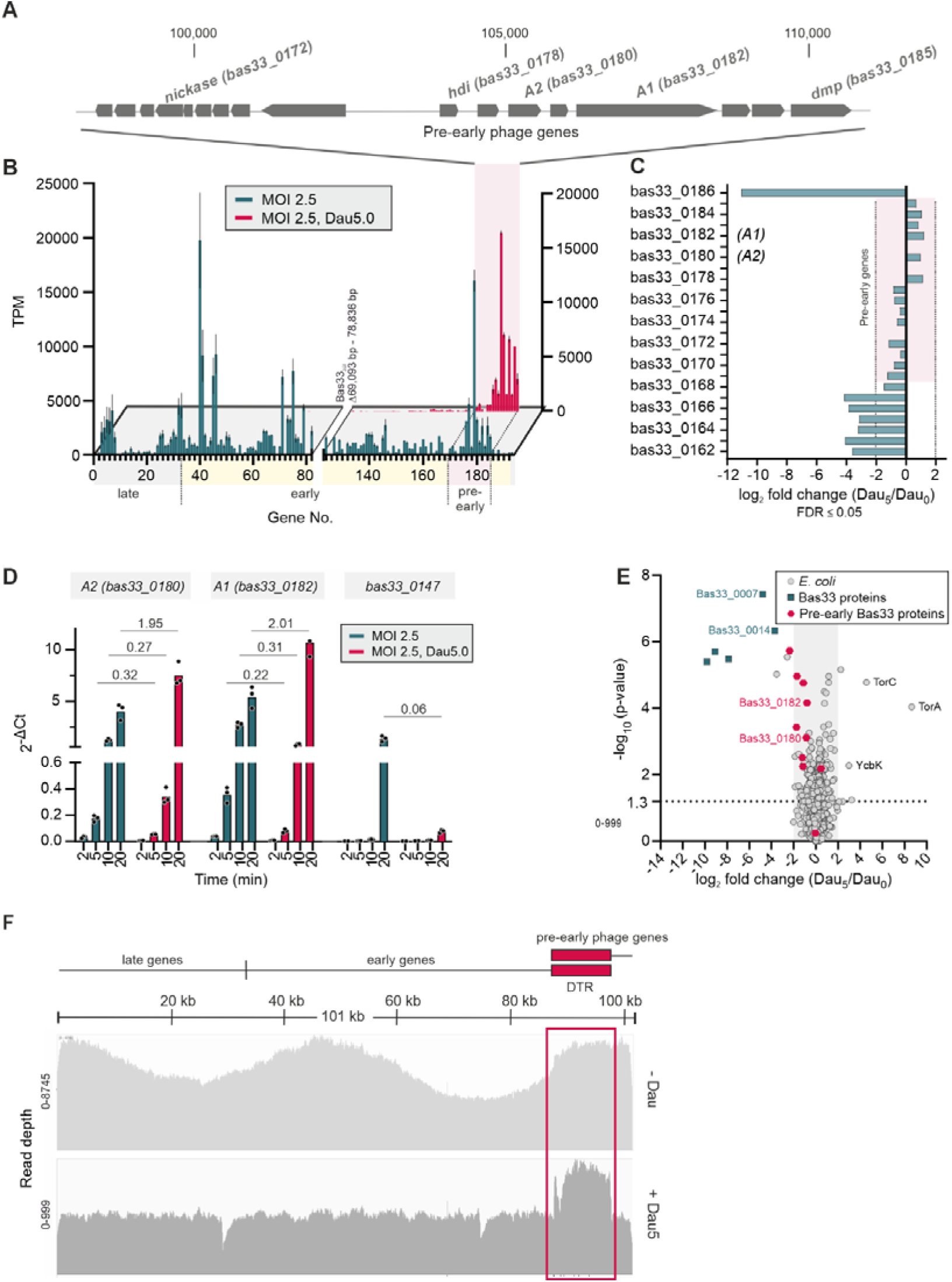
Transcriptomics and proteomics revealed a blockage of Bas33 phage gene expression after first-step transfer (FST). A) Pre-early gene region (8%) of the Bas33 genome (T5-like) with annotation of genes of known function. B) Mean values of detected phage transcripts (TPM) after 20 min of Bas33 infection using the E. coli K-12 MG1655 ΔRM as host background, shown as transcripts per million (TPM). Categorization into pre-early, early and late phage genes is highlighted alongside the gene numbers (n = 2). C) Zoom-in into log_2_ fold changes (Dau_5_/Dau_0_) of the pre-early gene region within the Bas33 genome showing no signifcant changes in transcription upon addition of daunorubicin. D) RT-qPCR showing time-resolved transcription of selected pre-early phage genes (Bas33_0180, coding for A2 protein, and Bas33_0182, coding for A1 protein) in comparison to an early phage gene (Bas33_0147, putative homing endonuclease)) in presence and absence of 5 µM daunorubicin. The relative copy number (GOI to housekeeping gene atpD of E. coli K-12 MG1655ΔRM) is shown as 2^-ΔCt^ for three independent biological replicates measured as technical duplicates. Mean values of calculated FC (Dau_5_/Dau_0_) at the different time points are indicated above. E) Volcano plot showing the proteome profile as log_2_ FC in protein abundance (Dau_5_/Dau_0_). Thresholds were set to log_2_ FC of 2 and -log_10_ (p-value) of 1.3 (moderated T-test, n = 3 biological replicates). F) Genom coverage after Oxford Nanopore sequencing of intracellular and cell-associated phage DNA of Bas33-infected E. coli K-12 MG1655 ΔRM (n=3). Predicted DTRs covering the pre-early phage gene region are indicated.

Remarkably, the host transcriptome of the *E. coli* K-12 MG1655 ΔRM strain exposed less differences with only 16 genes being upregulated (log_2_ FC ≥ 2, FDR ≤ 0.05) and 53 genes being downregulated (log_2_ FC ≤ -2, FDR ≤ 0.05; Table S1). Among the upregulated genes, the highest log_2_ FC (> 4.0) was detected for parts of the anaerobic respiratory chain (*torAC*) and three transposase genes (*insH1| insH10 | insL2*) as well as the putative fimbrial protein-encoding gene *ybgD*. The observed transcriptional pattern was consistent with the proteomic data, showing a similar protein abundance for the pre-early gene products. Without daunorubicin treatment, we reliably detected 122 Bas33 phage proteins. In contrast, in the presence of daunorubicin, only 15 phage proteins were reproducibly detected, 11 of which corresponded to pre-early gene products (Figure 3E).

Furthermore, we examined phage DNA replication using long-read sequencing (Figure 3F). When reads were mapped to the Bas33 genome, coverage was markedly higher under normal infection conditions, consistent with ongoing bidirectional replication, likely initiated from two origins. In contrast, in the presence of daunorubicin, coverage remained low and largely uniform across the genome, indicating that replication does not proceed. Notably, the pre-early phage gene region showed approximately twofold higher coverage relative to the rest of the genome, with sharp coverage drops at both boundaries. This pattern is readily explained by the presence of two copies of this region in the Bas33 genome, corresponding to the direct terminal repeat, as described for the model phage T5 (Davison, 2015, 2020). Additionally, the two distinct drops in the coverage plot may reflect single-stranded DNA (ssDNA) nicks, which are introduced into the 3L-5L strand of T5 genomes prior to encapsidation (Davison, 2015). This finding further supports the conclusion that phage DNA replication is blocked in the presence of daunorubicin.

### Synergy between daunorubicin-mediated defense and nucleic acid-targeting defense systems

Since DNA-intercalating compounds block phage infection at an early stage, we hypothesized that they could act synergistically with downstream DNA-targeting defense systems. Such systems have also been shown to primarily attack the pre-early phage gene region of T5-like phages (Ramirez-Chamorro et al., 2021; Strotskaya et al., 2017). To test this hypothesis, we selected a representative set of RM systems and evaluated their activity in combination with daunorubicin in a defined strain background. To this end, respective systems – showing either no, intermediate or strong defense against respective phages (Maffei et al., 2021) – were expressed from a low-copy plasmid under the native promotor in the *E. coli* K-12 MG1655 ΔRM strain, and infections were performed at low and high MOIs by initially focusing on *Demerecviridae*. However, members of the *Tequintavirus* genus, including phage Bas33, are largely resistant to RM systems, therefore we employed phage Bas29, a representative of the *Epseptimavirus* genus featuring sensitivity to RM type II systems EcoRV, and intermediate sensitivity to the type II and type III RM system EcoRI and EcoP1_I, respectively (Maffei et al., 2021). As expected, due to the absence recognition sites within the pre-early phage region, expression of EcoRI showed only minor difference compared to the growth control and retained the Abi-like phenotype at high MOIs and 2.5 µM daunorubicin. An intermediate effect was observed for the type II RM system EcoRV, preventing indeed culture collapse in absence of daunorubicin for low MOIs, but showing no strong effect on both conditions at high MOIs. The most pronounced effect was observed for EcoP1_I, preventing the growth arrest at high MOIs in presence of daunorubicin via synergistic interaction with daunorubicin as checked according to Wu et al. (2024) (Figure 4A and B). This is assumed to be conferred through the presence of 8 recognition sites already in the pre-early phage gene region.

**Figure 4:**
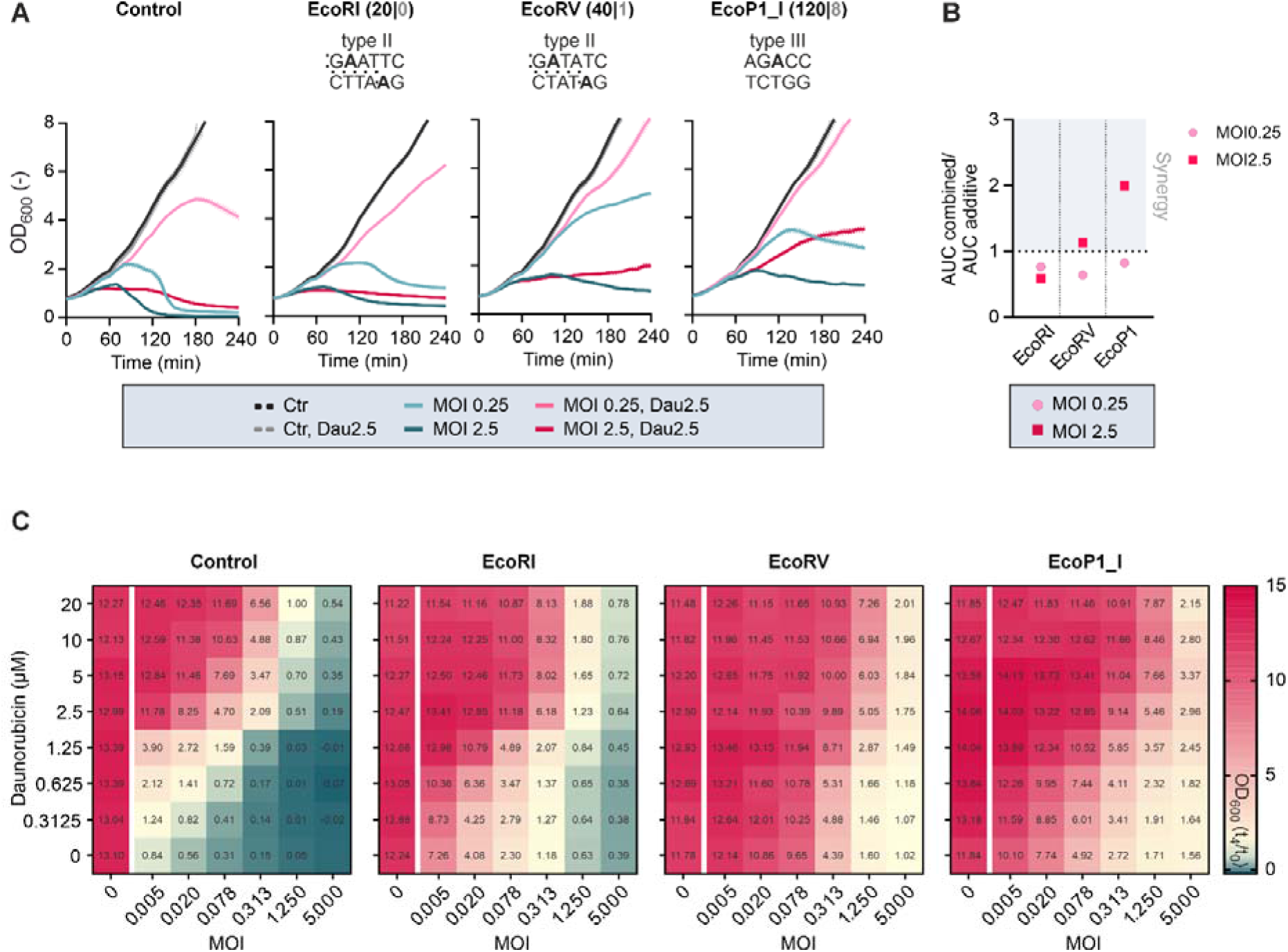
DNA-targeting defense systems show synergistic effects with daunorubicin. A) Liquid infection assays of the phage Bas29 infecting E. coli MG1655 ΔRM strains carrying different RM systems (control = empty plasmid). Assays were performed in presence and absence of 2.5 µM daunorubicin at low and high MOIs of 0.25 and 2.5 (n=3). Numbers of recognition sites of the respective RM systems in the entire genome and the pre-early gene region are indicated in black and gray, respectively. B) Calculation of synergy between daunorubicin and the respective ResMod systems based on the „Area under the curve (AUC)“ according to Wu et al. (2024) using OD_600_ recorded at t_0_ as baseline. Ratios were calculated from AUC mean values of three biological replicates. C) Checkerboard-like assays combining different MOIs of the phage Bas29 and different daunorubicin concentrations in the respective E. coli strains with distinct RM antiphage defense backgrounds. The panel shows the FC in OD_600_ after 4 h of cultivation and infection, with the calculated values indicated in the heatmaps.

To assess the synergistic effects more broadly, checkerboard-like assays combining different MOIs (0.005 to 5.0) with daunorubicin concentrations ranging from 0.3 to 20 µM were performed and FC in OD_600_ after 4 h was calculated to assess infection outcomes (Figure 4C). Control assays with the *E. coli* K-12 MG1655 ΔRM strain demonstrated that high MOIs combined with low daunorubicin concentrations promoted phage-induced cell lysis, whereas low MOIs and elevated daunorubicin concentrations favoured host survival. EcoRI expression caused a shift of the infection profile toward higher MOIs enabling cell survival up to an MOI of 0.3 in presence of ≥ 2.5 µM daunorubicin, and resulting in a less pronounced cell lysis at higher MOIs. EcoRV and EcoP1_I showed comparable infection dynamics, with cell survival observed at MOIs < 0.3 under all daunorubicin conditions, which was slightly more pronounced for EcoRV. Remarkably, the abi-like phenotype of the control and during EcoRI expression at MOIs of 1.25 was suppressed at daunorubicin concentrations of ≥ 2.5 µM.

Since T5-like phages are generally more resistant to defense via RM systems, we also analysed synergistic interactions of daunorubicin and RM systems during infection with the T1-like daunorubicin-sensitive phage Bas09 which shows high sensitivity to type II and III RM systems but resistance to the type I RM system EcoCFT_I. As expected due to the absence of multiple recognition sites, expression of EcoCFT_I exhibited the same phenotype as the empty plasmid control showing the Abi phenotype at high MOIs and 2.5 µM daunorubicin. An intermediate effect was observed for the type III RM system EcoCFT_II, leading to a delayed culture collapse at a low MOI in absence and no more growth defect in presence of daunorubicin. The most pronounced effect was observed for EcoRI, which strongly retarded the phage-mediated cell lysis under normal infection conditions and completely prevented the culture collapse and Abi phenotype for both MOIs upon addition of daunorubicin (Figure S4A). Checking for additive or synergistic effects revealed a synergistic interaction of EcoCFT_II and EcoRI with daunorubicin for both applied MOIs (Figure S4B). Checkerboard-like assays with Bas09 and the chosen RM systems further supported the synergistic action (Figure S4C).

Taken together, daunorubicin and restriction-modification systems can synergistically protect cells from phage-induced cell lysis or death, thereby allowing survival of the entire population. These results emphasize the intricate interaction between different lines and highlight the context-dependency of defense phenotypes.

## Discussion

Recent years have witnessed the discovery of numerous novel antiphage defense systems in bacteria, profoundly expanding our understanding of bacterial immunity. Beyond RNA or protein based systems also defense based on the activity of small antiphage molecules emerged as an underappreciated layer of phage defense (Hardy et al., 2023; Kever et al., 2022; Kronheim et al., 2018; Shomar et al., 2024). Among the most prominent examples are DNA-intercalating anthracyclines produced and secreted by *Streptomyces*, which were previously shown to block infection of various dsDNA phages at an early step of the phage life cycle (Kronheim et al., 2018). In this study, we used the *E. coli* BASEL phage collection (Maffei et al., 2021) to dissect the mechanism of action of DNA-intercalating antiphage molecules, focusing specifically on the anthracycline daunorubicin. While earlier work demonstrated a broad inhibitory effect, our systematic screening revealed pronounced phage-specific differences in phage sensitivity, with members of the *Drexlerviridae* (Bas1-13), *Demerecviridae* (Bas18-34) and *Vequinaviruses* (Bas48-62) showing particularly high sensitivity to DNA-intercalating antiphage compounds. Using the immunocompromised *E. coli* K-12 ΔRM strain (Maffei et al., 2021), we observed an ‘abortive infection’ (abi) phenotype for daunorubicin-sensitive phages (Aframian & Eldar, 2023; Lopatina et al., 2020). This was not only evidenced by applying high MOIs leading to the typical ‘abi-like’ culture collapse or growth stagnation, but was also shown via live cell imaging in microfluidic chip devices, confirming growth arrest followed by membrane permeabilization of infected cells in the presence of daunorubicin (Figure 2).

In our screening, members of the *Tevenvirinae*, including the model phage T4, exhibited a remarkably high resistance to different DNA-intercalating antiphage molecules (Figure 2F). Although phage epigenomic modifications are well known to protect phages from diverse nucleic acid-targeting defense systems (Weigele & Raleigh, 2016), our initial hypothesis, that the enhanced daunorubicin resistance of T4-like phages might result from DNA hypermodification, was disproved.

Using the T5-like phage Bas33, we observed that in the presence of daunorubicin transcription and translation were confined to the pre-early phage gene region injected during the first-step transfer (FST) (Davison, 2015). Furthermore, no phage DNA replication could be detected under daunorubicin pressure. Accordingly, we hypothesize that daunorubicin blocks Bas33 infection at FST level, either by preventing the transcription and replication of phage DNA regions injected secondly, or by directly preventing SST, which would result in only the first ∼9% of the Bas33 genome being injected into the host cell.

The molecular mechanism underlying SST still remains poorly understood. The injection stop signal (ISS) has been proposed to reside at the phage-host interface and is characterized by multiple repeats, inverted repeats, palindromic motifs and DnaA boxes. These features are thought to facilitate the formation of secondary structures and/or interaction with host components physically blocking SST (Davison, 2015, 2020; Heusterspreute et al., 1987). The phage proteins A1 ad A2, which are essential for SST, may involve DNA releasing the blockage, which in turn might be blocked by the DNA intercalating activity of daunorubicin. The second step transfer typically results in the repression of pre-early phage genes due to a shift in RNA polymerase specificity to early phage gene upon association with A1, which was absent under daunorubicin pressure. The deoxyribonucleotide 5L-monophosphatase (Dmp) promotes infection by further degrading host nucleotides during takeover and promoting their incorporation into viral DNA (Davison, 2015) (Davison, 2015). Additional toxicity to the host arises from the host division inhibitor (Hdi), which disrupts FtsZ ring formation, and the Ung-dependent nickase, which exploits Ung to target and cleave dUMP-containing DNA (Mahata et al., 2021; Mahata et al., 2023). Hence, the continued expression of the pre-early host takeover genes is likely to have toxic effects on the host cell, leading to cell death (Davison, 2015; Mahata et al., 2021; Mahata et al., 2023).

Similar to the pre-early proteins A1 and A2 of phage T5, many early phage proteins harm their host as an intrinsic part of the infection process (Davison, 2015; Souther et al., 1972; Svenson & Karlström, 1976). Accordingly, the accumulation of these (pre-)early phage proteins likely underlies the observed ‘abi’ phenotype. Importantly, in this case, cell death is not a result of a defense mechanism directed against host components, but rather an indirect effect from targeting phage components – defined as ‘mutual destruction’ (Aframian & Eldar, 2023). A similar phenotype was observed for CRISPR-Cas targeting the pre-early genes of T5, which also enforced abortive infection (Strotskaya et al., 2017). The distinction between classical abi and ‘mutual destruction’ caused by toxic phage products underscores the critical need to disentangle mechanistic cause from phenotypic outcome when interpreting abortive infection-like responses.

We furthermore show that the respective defense outcome is context dependent. While we do observe cell death in an immunocompromised strain background, the synergistic interaction of daunorubicin and nucleic-acid targeting defenses (RM systems EcoRV and EcoP1, Figure 4) counteracted the abi phenotype and enabled survival of the infected cells. This effect was dependent on the respective recognition sites within the pre-early phage gene region. Under these conditions, an assumed combined action of SST inhibition by daunorubicin and restriction of FST regions by RM systems probably prevents the accumulation of toxic phage products, thereby enabling cellular recovery. These findings align with recent studies emphasizing the context dependency of phage defense mechanisms (Wu et al., 2024). Another example involves type VI CRISPR systems, in which Cas13-mediated cleavage of mRNA targets would typically induce dormancy. However, in strains additionally carrying a RM system, initial dormancy is followed by recovery and resumed cellular growth (Labrie et al., 2010). This context-dependency even extends to the action of antibiotics as shown for the bactericidal activity of antifolate antibiotics in the presence of the CBASS antiphage defense system in *Vibrio cholera* (Brenzinger et al., 2024). Taken together, these studies highlight the complex interplay between small molecules and the different layers of the bacterial immune system, underscoring how the genetic makeup of the host strain shapes the resulting defense phenotype.

## Material and methods

### Bacterial strains and infection conditions

All bacterial strains, phages and plasmids used in this study are listed in Table S2, S3 and S4, respectively. During this study, we observed that our variant of Bas33 carries a deletion (Bas33 Δ69,093 - 78,836 bp; Figure S5) at a locus where similar deletions had also been previously observed for variants of its close relative T5 (Burman et al., 2024). Since phenotypes observed with Bas33 were consistent with those for other *Markadamsvirinae* phages (Figure 1, 2) this deletion does not affect the phenotypes described in this manuscript.

*Escherichia coli* cultures were inoculated from single colonies grown on LB agar plates in 5 ml LB medium supplemented with the indicated antibiotics and cultivated for 16 h at 37°C and 170 rpm. For infection assays in liquid cultures, 300 µl main cultures were inoculated from pre-cultures in the same medium to the desired OD_600_ and cultivated GrowthProfiler microcultivation system (Enzyscreen, Heemstede, The Netherlands) at 225 rpm and 37°C using a 10 min imaging interval. Daunorubicin was directly added to the main cultures to the indicated concentration. Red values were recorded and converted to OD via calibration curves determined via the strain calibration data sheet of (Enzyscreen, Heemstede, The Netherlands):

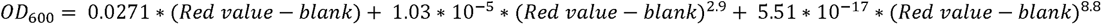

For omics-approaches, cultivations were scaled up to shaking flasks and cultivation was done at 150 rpm and 37°C. Infection with phages at the indicated MOI was done after inoculation and daunorubicin addition with a 5 minute delay. If not indicated otherwise, infection assays were done in three independent biological replicates.

For double-agar overlay assays, *E. coli* K-12 MG1655 ΔRM pre-cultures were used to inoculate LB soft-agar (0.4%) to an OD_600_ of 0.2 and poured on LB agar plates. Decimal dilution series of phages in SM buffer (50 mM Tris-HCl, 100 mM NaCl, 8 mM MgSO_4_, pH 7.5) were spotted on top. Daunorubicin (dissolved in DMSO) was added to both agar layers to the indicated concentrations.

### Phage amplification and purification

For phage amplification, 5 ml *E. coli* K-12 MG1655 ΔRM pre-cultures in LB medium were used to inoculate 70 ml main cultures in the same medium to a starting OD_600_ of 0.2. Cells were cultivated at 37°C and 120 rpm for 90 min before phages were added to an MOI of 0.01. Supernatant was harvested after culture clearance via centrifugation at 5,000 *g* and 40°C for 10 min. After sterile filtration, the phage lysate was again centrifuged at 37,000 *g* and 4°C for 90 min. The phage pellet was resuspended in 1 ml TM buffer (50 mM Tris-HCl, 10 mM MgCl_2_, pH 7.5) and stacked on top of a sucrose gradient (0-45%; in TM buffer). Ultracentrifugation was performed at 70,000 *g* and 4°C for 20 min. The phage band was extracted using a syringe with needle and transferred to a fresh ultracentrifugation tube. 4 ml SM buffer were added and centrifugation was repeated at 70,000 *g* and 4°C for 90 min. The phage pellet was finally resuspended in 4 ml SM buffer, sterile filtrated and titers were determined via double-agar overlay assays.

### Live cell imaging on single-cell level in microfluidic devices

Single-cell analysis of infection dynamics was performed using an in-house developed microfluidic platform (Grünberger et al., 2015; Grünberger et al., 2013). Cultivation of *E. coli* K-12 MG1655 ΔRM was performed in 60×60 µM chambers (height: 1020-1040 nM) flushed with LB medium containing 20 µM propidium iodide and 2.5 µM daunorubicin if indicated. A concentration of 10^9^ PFU/ml phages was continuously added with the medium flow at a flow rate of 200 nl min^-1^. Growth and cell fate was recorded in 5 min intervals using phase contrast and the optical filter TexasRed (excitation, 560/40 nm; dichroic, 595; emission, 630/60 nm) at an exposure time of 200 ms and 100 ms, respectively. Preparation of image cutouts and adjustments of lookup tables (LUTs) were performed using NIS-Elements BR 5.30.03 (64 bit) and ImageJ 1.54f (Rueden et al., 2017). Single-cell analysis has been performed based on acia-workflows (Seiffarth et al., 2025) with custom single-cell analysis. The amount of lysed cells is determined by (1) segmenting cells using Omnipose (Cutler et al., 2022), (2) cell size based filtering, and (3) computing the temporal cell population development (based on count and area) averaged over triplicates. The 50% drop from the global maximum quantifies the cell lysing progress. To determine single-cell PI fluorescence readouts, cell segments are tracked using trackastra (Gallusser & Weigert, 2025) and the fluorescence development for every individual cells is measured. The developed acia-workflows performing these analysis steps are provided with the supplementary information.

### Quantification of daunorubicin uptake

For measurement of daunorubicin uptake upon phage infection, *E. coli* K-12 MG1655 ΔRM was cultivated as described in ‘Bacterial strains and infection conditions’ using an initial OD_600_ of 1.0 and an MOI of 2.5. Cultivation was performed in 1 ml Eppendorf tubes at 37°C and 500 rpm. At the indicated time points, 250 µl cells were harvested by centrifugation at 16,000 *g* and 4°C for 2 min and subsequently resuspended in 250 µl PBS. For measurement, 2 µl cells were diluted in 1 ml dd H_2_O and daunorubicin uptake was quantified via fluorescence measurements with the Cytek® Aurora flow cytometer using the detectors of the 488 nm laser. The gate was set up to gate on bacteria in FSC/SSC, followed by doublet exclusion by FSC-A/FSC-H. A gate for positive events was set based on unstained controls capturing 0.1% of the population at linear gate. Readouts were done for the percentage positive cells as well as for the MFI (mean of fluorescence) of the entire singlet population to monitor the shift in signal intensities.

### Long-read sequencing of phage-bacteria complexes

*E. coli* K-12 MG1655 ΔRM was cultivated as described in ‘Bacterial strains and infection conditions’ using an initial OD_600_ 0.8 and MOI of 2.5 in a total volume of 10 ml LB medium ± 2.5 µM daunorubicin. After 20 min of infection, cultures were harvested by centrifugation at 5,000 *g* and 4°C for 10 min and washed two times with 10 ml 0.9% NaCl to remove unbound phages. For DNA isolation, the cells were resuspended in 1 ml Tris buffer (20 mM Tris-HCl, 300 mM NaCl, pH 8.0), transferred to cell disruption tubes containing glass beads, and disrupted twice at 6,500 *g* for 20 sec using a Precellys® cell disrupter. After centrifugation at 16,000 *g* and 4 °C for 5 min, supernatants were transferred to a fresh tubes and RNA and protein digestion was performed with 200 µg/ml RNase A for 30 min at 30°C and 300 µg/ml proteinase K for 2 h at 55°C, respectively. Finally, the bacterial and phage DNA was purified using three cycles of phenol/chloroform/isoamylalcohol extraction and three cycles of chloroform extraction. The DNA was then precipitated using 0.1 volumes of 3 M sodium acetate (CH_3_COONa) and 2.5 volumes of absolute ethanol at - 20°C for 16 h, dried and resuspended in 50 µl dd.H_2_O. Total DNA was subjected to Oxford Nanopore sequencing using the Native Barcoding Kit (SQK-NBD114.24) on an R10.4.1 flow cell (FLO-PRO114M) following the manufacturer’s instructions (Oxford Nanopore Technologies, UK).

Raw sequencing data were basecalled with Dorado version 1.0.2 (Oxford Nanopore Technologies, UK) using the „super“ accurate model dna_r10.4.1_e8.2_400bps_sup@v5.2.0 and the parameter --kit-name SQK-NBD114-24. Demultiplexing was performed with Dorado demux (version 1.0.2) using the --no-classify option. Reads from each replicate and condition were aligned against a combined reference consisting of *E. coli* K-12 MG1655 (NCBI:NC_000913.3) and *Escherichia* phage HildyBeyeler (Bas33; MZ501074.1). Alignments were generated with minimap2 (version 2.30) using the presets -ax map-ont and the option --secondary=no (Li, 2018). Non-primary alignments, duplicates, and supplementary alignments were excluded with samtools view (version 1.20) using --excl-flags 3328 and converted to bam format with –bam (Danecek et al., 2021). Reads mapping exclusively to the *Escherichia* phage HildyBeyeler (*Bas33*) reference were extracted with samtools fastq (version 1.20) with default settings (Danecek et al., 2021). Subsequent filtering using seqkit (version 2.9.0) retained only reads with a minimum quality score of 20 and a length of at least 10 kb (Shen et al., 2024). Assembly of filtered reads was conducted with hifiasm (version 0.25.0) using the ont parameter (Cheng et al., 2021). The resulting contig was linearized by identifying the terminase small subunit (*terS*) as the starting position, in accordance with the Bas33 reference genome.

Annotation of the hifiasm assembly was performed with Prodigal (version 2.6.3) to predict protein-coding sequences (Hyatt et al., 2010). Putative *terL* and *terS* proteins were identified by searching the UniProt database (UniProt Consortium 2017). Reviewed sequences were used to construct two BLAST protein databases with makeblastdb (version 2.15.0) using the options -dbtype prot and -parse_seqids. Searches were then performed with blastp (version 2.15.0) using the parameters -evalue 1e-5, -max_target_seqs 10, and -outfmt 6 (Altschul et al., 1990; Camacho et al., 2009). The identified *terS* hit was then used to confirm the start position for contig linearization. For comparative alignment, the re-assembled Bas33 genome from this study was combined with the *E. coli* K-12 MG1655 reference and re-analyzed using the same mapping and filtering workflow as described above. Only alignments to the assembled phage reference were retained and visualized with Integrative Genomics Viewer (IGV) (version 2.16.2) (Robinson et al., 2011).

### Transcriptomics via RNA sequencing

For transcriptomic analyses, cells were cultivated and infected as described in ‘Bacterial strains and infection conditions’ using an initial OD_600_ of 0.8 and an MOI of 2.5 in a total volume of 25 ml LB medium, supplemented with 5 µM daunorubicin if indicated. At 20 min post infection, cells were harvested on ice at 5,000 g and 4°C for 7 min and cell pellets were frozen in liquid nitrogen. Subsequent RNA isolation was done with the Monarch Total RNA Miniprep Kit (NEB, Massachusetts, US) and concentrations were determined using NanoDrop Spectrophotometer. RNA depletion, library preparation and paired-end sequencing was performed at GENEWIZ from Azenta Life Sciences (Leipzig, Germany). RNA sequencing results were analysed using the CLC genomics workbench v.20 (Qiagen, Germany) as described in Luthe et al. (2023).

### Transcriptomics via RT-qPCR

For quantification of single gene transcription of the Bas33 phage, cultivation, cell harvesting and RNA isolation was performed as described in, Annotation of the hifiasm assembly was performed with Prodigal (version 2.6.3) to predict protein-coding sequences (Hyatt et al., 2010). Putative *terL* and *terS* proteins were identified by searching the UniProt database (UniProt Consortium 2017). Reviewed sequences were used to construct two BLAST protein databases with makeblastdb (version 2.15.0) using the options -dbtype prot and -parse_seqids. Searches were then performed with blastp (version 2.15.0) using the parameters - evalue 1e-5, -max_target_seqs 10, and -outfmt 6 (Altschul et al., 1990; Camacho et al., 2009). The identified *terS* hit was then used to confirm the start position for contig linearization. For comparative alignment, the re-assembled Bas33 genome from this study was combined with the *E. coli* K-12 MG1655 reference and re-analyzed using the same mapping and filtering workflow as described above. Only alignments to the assembled phage reference were retained and visualized with Integrative Genomics Viewer (IGV) (version 2.16.2) (Robinson et al., 2011).

Transcriptomics via RNA sequencing‘ at the indicated time points. RT-qPCR was performed with the Luna One-Step RT-qPCR Kit (New England BioLabs, Frankfurt am Main) according to the manufacturer’s instructions in the qTower (Analytik Jena, Jena) using an input concentration of 25 ng/µl total RNA. Transcripts of *bas33_0180* (coding for A2 protein), *bas33_0182* (coding for A1 protein and *bas33_0147* (coding for an homing endonuclease) were quantified using the *atpD* of *E. coli* K-12 MG1655 ΔRM as reference gene. Data were analyzed with qPCRsoft 3.1 (Analytik Jena, Jena) and relative concentrations were calculated according to the 2^−ΔCt^ method (Livak & Schmittgen, 2001). Primers used for amplification are provided in Table S5.

### Proteomics via LC-MS

For proteome analysis, the biological triplicates were taken from the same cultivation batch as for the transcriptomic analyses by harvesting 2 ml cells at 16,000 g and 4°C for 2 min and subsequent freezing of cell pellets in liquid nitrogen.

In order to prepare samples for proteome measurements, cells were thawed on ice and lysed by resuspending in 200 µL of lysis buffer (2% sodium lauroyl sarcosinate (SLS) in 100 mM ammonium bicarbonate (ABC) buffer), heating at 90°C for 15 min and 3 cycles of sonication (30s each) using a Hielscher Up200St ultrasonic processor (75% amplitude, 0.5 s / 0.5 s pulse). Subsequently, samples were centrifuged at 14,000 rpm for 5 min and supernatants were transferred into a new low protein binding tube. Protein concentrations were determined using a Pierce^TM^ BCA Protein assay kit (ThermoFisher Scientific, 23250). Afterwards, 2 mM final <u>tri</u>s(2-carboxyethyl)phosphine (TCEP, Tokyo Chemical Industry, ref. T1656) were added and samples were mixed and incubated at 90°C for 15 min to promote reduction of disulfide bridges. After cooling down, 4 mM iodoacetamide were added and samples were incubated in the dark for 30 min at room temperature. Afterwards, 30 µg of total lysate proteins were transferred to a new low protein binding tube and sample volume was adjusted with lysis buffer to a total volume of 50 µL. Then, 3 volumes of 100 mM ABC buffer were added to the sample in order to dilute the final SLS concentration down to 0.5%. Then, SP3 magnetic beads slurry was prepared as follow: in a new tube, 20 µL of beads (Sera-Mag^TM^ carboxylate-modified [E3] and [E7] magnetic beads, Cytiva) were mixed at a 1:1 ratio and washed twice with 300 µL of HPLC grade water. Then, beads were resuspended in 100 µL of water to produce readily usable slurry. Afterwards, 8 µL of bead slurry was added to the protein sample along with 3 sample volumes (600 µl) of 100% acetonitrile (ACN). Proteins were allowed to bind to the beads for 1 h at RT with 800 rpm shaking. Supernatant was discarded by using a magentic rack and beads were washed twice with 300 µl of 70% ethanol, once with 300 µL of 100% ACN before air drying for 5 min. Finally, protein digest was performed by adding 200 µL of 100 mM ABC buffer and 0.6 µg of sequencing-grade trypsin (Promega, V5111) overnight at 30°C with 1,000 rpm shaking. Supernatants were transferred to a new low protein binding tube and beads were resuspended in 100 µL of Millipore H_2_O. After magnetic bead separation, supernatants were pooled with the previous ones and residual SLS was precipitated by addition of 4% final concentration of trifluoroacetic acid (TFA, Serva 45641.01). Samples were centrifuged at 4°C at 14,000 rpm for 20 min and supernatants were purified with C18 columns (Chromabond, 730522.250). The solvent was evaporated with SpeedVac vacuum concentrations at 45°C and stored samples at -20°C before measurement.

Liquid Chromatography-Mass Spectrometry (LC-MS) for peptide analysis was carried out using a Vanquish Neo system coupled to an Orbitrap Exploris 480 mass spectrometer. Peptides were dissolved 0.1% trifluoroacetic acid and loaded onto a self-packed C18 (26 cm of 1.9 μm Reprosil-AQ, Dr. Maisch) column. The peptides were separated by an acetonitrile gradient running from 2 – 25 % solvent B (99.85% acetonitrile, 0.15% formic acid) over 45 min, followed by an additional increase of solvent B to 35% and 40 %, respectively, for 15 min at a flow rate of 300 nl/min. Solvent A composition was 0.15 % formic acid. Eluting peptides were analyzed in data independent acquisition (DIA) mode on the Exploris-MS. The funnel RF level was set to 40. Full MS resolution was set to 120.000 (m/z 200). Automatic gain control (AGC) target value for fragment spectra was set at 3000%. 45 windows of 14 Da were used with an overlap of 1 Da between m/z 320-950. Resolution was set to 15,000 and fill time to 22 ms. Stepped HCD collision energy of 25, 27.5, 30 % was used. MS1 data was acquired in profile, MS2 DIA data in centroid mode.

Analysis of DIA data was performed using the DIA-NN version 1.8 and 1.9 (Demichev et al., 2020), respectively, using a uniprot protein database from E.coli with Bas33 phage proteins included (MZ501074; UniProt ID: 2852005) to generate a data set specific spectral library for the DIA analysis. The neural network based DIA-NN suite performed noise interference correction (mass correction, RT prediction and precursor/fragment co-elution correlation) and peptide precursor signal extraction of the DIA-NN raw data. The following parameters were used: Full tryptic digest was allowed with two missed cleavage sites, and oxidized methionines (variable) and carbamidomethylated cysteins (fixed). Match between runs and remove likely interferences were enabled. The precursor FDR was set to 1%. The neural network classifier was set to the single-pass mode. Quantification strategy was set to any LC (“high accuracy” for DIA-NN 1.8) and Quant UMS (“high precision” for DIA-NN 1.9). Cross-run normalization was set to RT-dependent. Library generation was set to smart profiling. DIA-NN outputs were further evaluated using the SafeQuant (Ahrné et al., 2013; Glatter et al., 2012) script modified to process DIA-NN outputs.

Data visualization was performed using GraphPad Prism 10.6.1. In brief, *E. coli* proteins that had a peptide count across replicates lower or equal to one count were filtered out and omitted from the analysis. Significance was calculated using a moderated t-test. The significance threshold for the -Log_10_(*p-value*) parameter was set to 1.3 and the threshold for the Log_2_(ratio) value was set to 2. Phage proteins were considered reliably detected if their average peptide count across replicates was ≥ 2.

## Supplemental information

Table S1 – S5

Figure S1 – S5

Videos S1 – S2

## Acknowledgments

We thank the Deutsche Forschungsgemeinschaft (SPP 2330 “New concepts in prokaryotic virus-host interaction”, project 464434020, SFB 1535 “Microbial Networking”, project ID 458090666) for financial support. K.H. is supported by funding from the Max Planck Society, the SPP 2330 (project ID 464500427), and RTG 2937 “Microbial Nucleotide Metabolism”. We thank Marianne De-Paepe for providing the T4 Δgt mutant phage. We furthermore thank Timo Glatter and Jörg Kahnt for the mass spectrometry and proteomics measurements.

## Author contributions

Conceptualization: LE | BR | JF

Data curation: LE | BR | JS | MB | EG | VR

Formal analysis: LE | BR | JS | MB | EG | NP |VR

Funding acquisition: JF | US | DK | KH

Investigation: LE | CG | BR | JS | MB | EG | VR

Methodology: LE | CG | JS | MB | EG | VR | NP | JF | KH

Resources: AH | NP | KH

Project administration: JF | LE

Supervision: JF | KH

Validation: LE | BR | JS | MB | EG | VR | NP

Visualization: LE | JS | MB | EG | VR

Writing – original draft: LE | JF

Writing – review & editing: all

## Data availability

The RNA-seq data were deposited in the European Nucleotide Archive (ENA) at EMBL-EBI (https://www.ebi.ac.uk/ena/browser/home) under accession number PRJEB101990. The long read sequencing data were deposited in the European Nucleotide Archive (ENA) at EMBL-EBI (https://www.ebi.ac.uk/ena/browser/home) under accession number PRJEB103991. The mass spectrometry proteomics data have been deposited to the ProteomeXchange Consortium via the PRIDE (Perez-Riverol et al., 2024) partner repository with the dataset identifier PXD070949.

## Declaration of generative AI and AI-assisted technologies in the writing process

During the preparation of this work the authors used ChatGPT in order to improve language of the manuscript. After using this tool, the authors reviewed and edited the content as needed and take full responsibility for the content of the published article.

