## Supplemental Information for "DNA-intercalating antiphage molecules trigger abortive infection through ‘mutual destruction’ and synergize with bacterial immunity"

#### Supplementary Tables

Table S1: Transcripts per million and differential expression upon Bas33 infection in presence and absence of 5 µM daunorubicin at 20 min post infection.

Table S2: Bacterial strains used in this study

Table S3: Phages used in this study

Table S4: Plasmids used in this study

Table S5: Oligonucleotides used in this study

#### Supplementary Figures

Figure S1: DNA hypermodification showed no influence on sensitivity to daunorubicin

Figure S2: Growth and fluorescence analysis on single-cell level during cultivation of E. coli K-12 MG1655 [illegible]RM in microfluidic chips with applied propidium iodide (PI) stain

Figure S3: Influence of different DNA-intercalating agents on phage infection dynamics

Figure S4: DNA-targeting defense systems showed synergistic effects with daunorubicin towards Bas09 infection

Figure S5: Comparison of Bas33$_{\text{wildtype}}$ and Bas33$_{\text{Jülich}}$ genome

#### Supplementary Videos

Video S1: Infection of *E. coli* K-12 MG1655 DRM with Bas33 in microfluidic chips using LB medium.

Video S2: Infection of *E. coli* K-12 MG1655 DRM with Bas33 in microfluidic chips using LB medium with 2.5 µM Daunorubicin.

### Supplementary Tables

**Table S1: Transcripts per million and differential expression upon Bas33 infection in presence and absence of 5 µM daunorubicin at 20 min post infection.**

This table is provided as separate file: Table 5_Expression Browser_TPM_DifExp_Bas33_20 min

**Table S2: Bacterial strains used in this study**

| Bacterial strains | Description | Reference |
|---|---|---|
| *Escherichia coli* K-12 MG1655 | $F^-$, $\lambda^-$, *ilvG*$^-$, *rfb-50 rph-1* | |
| *Escherichia coli* K-12 MG1655 ΔRM | *E. coli* K-12 MG1655 *Δmrr-hsdRMS-mcrBC* Δ*mcrA* | (Maffei et al., 2021) |
| *Escherichia coli* K-12 MG1655 ΔRM_pBR322_ΔP*tet* | *E. coli* K12 MG1655 ΔRM carrying the plasmid pBR322_Δ$P_{tet}$, $Amp^R$ | This work |
| *Escherichia coli* K-12 MG1655 ΔRM_pEcoRI | *E. coli* K12 MG1655 ΔRM carrying the plasmid pEcoRI, $Amp^R$ | This work |
| *Escherichia coli* K-12 MG1655 ΔRM_pAH213_EcoCFT_I | *E. coli* K12 MG1655 ΔRM carrying the plasmid pAH213_EcoCFT_I, $Amp^R$ | This work |
| *Escherichia coli* K-12 MG1655 ΔRM_pAH213_EcoECF_II | *E. coli* K12 MG1655 ΔRM carrying the plasmid pAH213_EcoCFT_II, $Amp^R$ | This work |
| *Escherichia coli* K-12 MG1655 ΔRM_pAH213_EcoP1_I | *E. coli* K12 MG1655 ΔRM carrying the plasmid pAH213_EcoP1_I, $Amp^R$ | This work |

**Table S31: Phages used in this study**

| Phage | Phage family | Subfamily | Bacterial host strain | Reference |
|---|---|---|---|---|
| Bas01 | *Drexlerviridae* | *Braunvirinae* | *E. coli* MG1655 $\Delta$RM | (Maffei et al., 2021) |
| Bas02 | *Drexlerviridae* | *Braunvirinae* | | |
| Bas03 | *Drexlerviridae* | *Braunvirinae* | | |
| Bas04 | *Drexlerviridae* | *Tempevirinae* | | |
| Bas05 | *Drexlerviridae* | *Tempevirinae* | | |
| Bas06 | *Drexlerviridae* | *Tempevirinae* | | |
| Bas07 | *Drexlerviridae* | *Tempevirinae* | | |
| Bas08 | *Drexlerviridae* | *Tempevirinae* | | |
| Bas09 | *Drexlerviridae* | *Tempevirinae* | | |
| Bas10 | *Drexlerviridae* | *Tempevirinae* | | |
| Bas11 | *Drexlerviridae* | *Tempevirinae* | | |
| Bas12 | *Drexlerviridae* | *Tunavirinae* | | |
| Bas13 | *Drexlerviridae* | *Tunavirinae* | | |
| Bas14 | | | | |
| Bas15 | | | | |
| Bas16 | | | | |
| Bas17 | | | | |
| Bas18 | | | | |
| Bas19 | | *Queuovirinae* | | |
| Bas20 | | *Queuovirinae* | | |
| Bas21 | | *Queuovirinae* | | |
| Bas22 | | *Queuovirinae* | | |
| Bas23 | | *Queuovirinae* | | |
| Bas24 | | *Queuovirinae* | | |
| Bas25 | | *Queuovirinae* | | |
| Bas26 | *Demerecviridae* | *Markadamsvirinae* | | |
| Bas27 | *Demerecviridae* | *Markadamsvirinae* | | |
| Bas28 | *Demerecviridae* | *Markadamsvirinae* | | |
| Bas29 | *Demerecviridae* | *Markadamsvirinae* | | |
| Bas30 | *Demerecviridae* | *Markadamsvirinae* | | |
| Bas31 | *Demerecviridae* | *Markadamsvirinae* | | |
| Bas32 | *Demerecviridae* | *Markadamsvirinae* | | |
| Bas33 | *Demerecviridae* | *Markadamsvirinae* | | |
| Bas34 | *Demerecviridae* | *Markadamsvirinae* | | |
| Bas35 | *Straboviridae* | *Tevenvirinae* | | |
| Bas36 | *Straboviridae* | *Tevenvirinae* | | |
| Bas37 | *Straboviridae* | *Tevenvirinae* | | |
| Bas38 | *Straboviridae* | *Tevenvirinae* | | |
| Bas39 | *Straboviridae* | *Tevenvirinae* | | |
| Bas40 | *Straboviridae* | *Tevenvirinae* | | |
| Bas41 | *Straboviridae* | *Tevenvirinae* | | |
| Bas42 | *Straboviridae* | *Tevenvirinae* | | |
| Bas43 | *Straboviridae* | *Tevenvirinae* | | |
| Bas44 | *Straboviridae* | *Tevenvirinae* | | |
| Bas45 | *Straboviridae* | *Tevenvirinae* | | |
| Bas46 | *Straboviridae* | *Tevenvirinae* | | |

| | | | | |
|---|---|---|---|---|
| Bas47 | *Straboviridae* | *Tevenvirinae* | | |
| Bas48 | | *Vequintavirinae* | | |
| Bas49 | | *Vequintavirinae* | | |
| Bas50 | | *Vequintavirinae* | | |
| Bas51 | | *Vequintavirinae* | | |
| Bas52 | | *Vequintavirinae* | | |
| Bas53 | | *Vequintavirinae* | | |
| Bas54 | | *Vequintavirinae* | | |
| Bas55 | | *Vequintavirinae* | | |
| Bas56 | | *Vequintavirinae* | | |
| Bas57 | | *Vequintavirinae* | | |
| Bas58 | | *Vequintavirinae* | | |
| Bas59 | | *Vequintavirinae* | | |
| Bas60 | | *Stephanstirmvirinae* | | |
| Bas61 | | *Stephanstirmvirinae* | | |
| Bas62 | | *Stephanstirmvirinae* | | |
| Bas63 | *Andersonviridae* | *Ounavirinae* | | |
| Bas64 | *Autographviridae* | *Studiervirinae* | | |
| Bas65 | *Autographviridae* | *Studiervirinae* | | |
| Bas66 | *Autographviridae* | *Studiervirinae* | | |
| Bas67 | *Autographviridae* | *Studiervirinae* | | |
| Bas68 | *Autographviridae* | *Studiervirinae* | | |
| Bas69 | *Schitoviridae* | *Enquatrovirinae* | | |
| $Bas33_{Jül}$ (Bas33 Δ69,093 - 78,836 bp) | *Demerecviridae* | *Markadamsvirinae* | *E. coli* MG1655 ΔRM | This study* |
| T4 | *Straboviridae* | *Tevenvirinae* | *E. coli* B (DSM613) | DSM 4505 |
| T5 | *Demerecviridae* | *Markadamsvirinae* | *E. coli* B (DSM613) | DSM 16353 |
| T7 | *Autographviridae* | *Studiervirinae* | *E. coli* B (DSM613) | DSM 4623 |
| Lambda | | | *E. coli* LE392 (DSM4230) | DSMZ |
| T4 Δα/β gt | *Straboviridae* | *Tevenvirinae* | *E. coli* B (DSM613) | gifted by Marianne De-Paepe |

*For all assays, the phage variant $Bas33_{Jül}$ was used (Figure S5).

**Table S4: Plasmids used in this study**

| Plasmid | Description | Reference |
|---|---|---|
| **pBR322_ΔP*tet*** | $Amp^R$, Derivative of pBR322, tetracycline resistance cassette deleted, empty vector control | (Pleška et al., 2016) |
| **pEcoRI** | $Amp^R$, derivative of pBR322 expressing *E. coli* type II RM system EcoRI | |
| **pEcoRV** | $Amp^R$, derivative of pBR322 expressing *E. coli* type II RM system EcoRV | |
| **pAH213_EcoCFT_I** | $Amp^R$, derivative of pBR322 expressing *E. coli* type I RM system EcoCFT_I of *E. coli* CFT073 | (Maffei et al., 2021) |
| **pAH213_EcoECF_II** | $Amp^R$, derivative of pBR322 expressing *E. coli* type III RM system EcoCFT_II of *E. coli* CFT073 | |
| **pAH213_EcoP1_I** | $Amp^R$, derivative of pBR322 expressing *E. coli* type III RM system EcoP1_I of *E. coli* phage P1 | |

**Table S5: Oligonucleotides used in this study**

| Name | Oligonucleotide sequence (5'-3') |
|---|---|
| **Bas33_qPCR_0180_fw** | GGCTTCTCCCGTGTCCGTTC |
| **Bas33_qPCR_0180_rv** | CCAGATCATCGGCGTCCTTGA |
| **Bas33_qPCR_0182_fw** | GCGGCGAAAGTAGCAAAAGCC |
| **Bas33_qPCR_0182_rv** | TATTGTTCAAGCGCGGCCAAAAAG |
| **Bas33_qPCR_0147_fw** | TCCCGAACAACGTGAAGATCGC |
| **Bas33_qPCR_0147_rv** | CGTGGCAATCTACAGCTTCCCAA |
| ***E. coli*_qPCR_*atpD*_fw** | ACGGAATTTCTCAGCCATGGTCAGAC |
| ***E. coli*_qPCR_*atpD*_rv** | GTGTTTGCGGGCGTAGGTGAAC |

#### Supplementary Figures

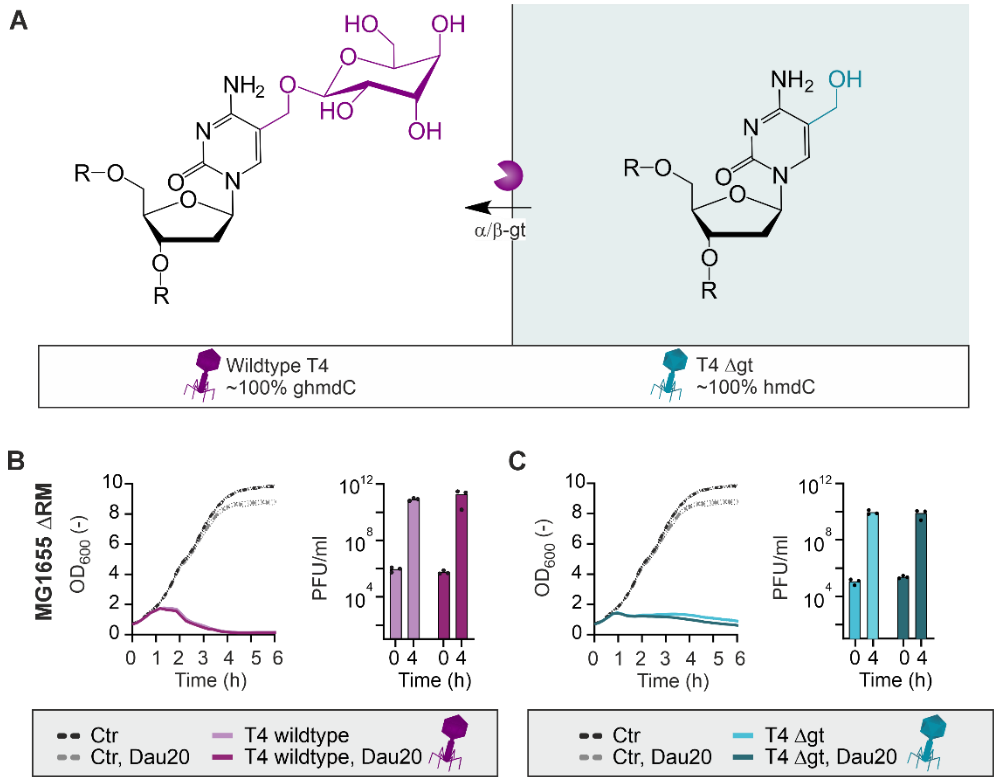


**Figure S1: DNA hypermodification showed no influence on sensitivity to daunorubicin.** A) Schematic representation of DNA modifications in wildtype T4 phages having glycosylated hydroxylmethyl-dCTPs (ghmdC) and T4 Δgt mutant phages lacking the glycosyl group (hmdC). B) Growth curves and titer development of *E. coli* K-12 MG1655 ΔRM upon infection with T4 phage in presence and absence of 20 µM daunorubicin. C) Growth curves and titer development of *E. coli* K-12 MG1655 ΔRM upon infection with T4 Δgt mutant phages in presence and absence of 20 µM daunorubicin. All assays were performed as three independent biological replicates.

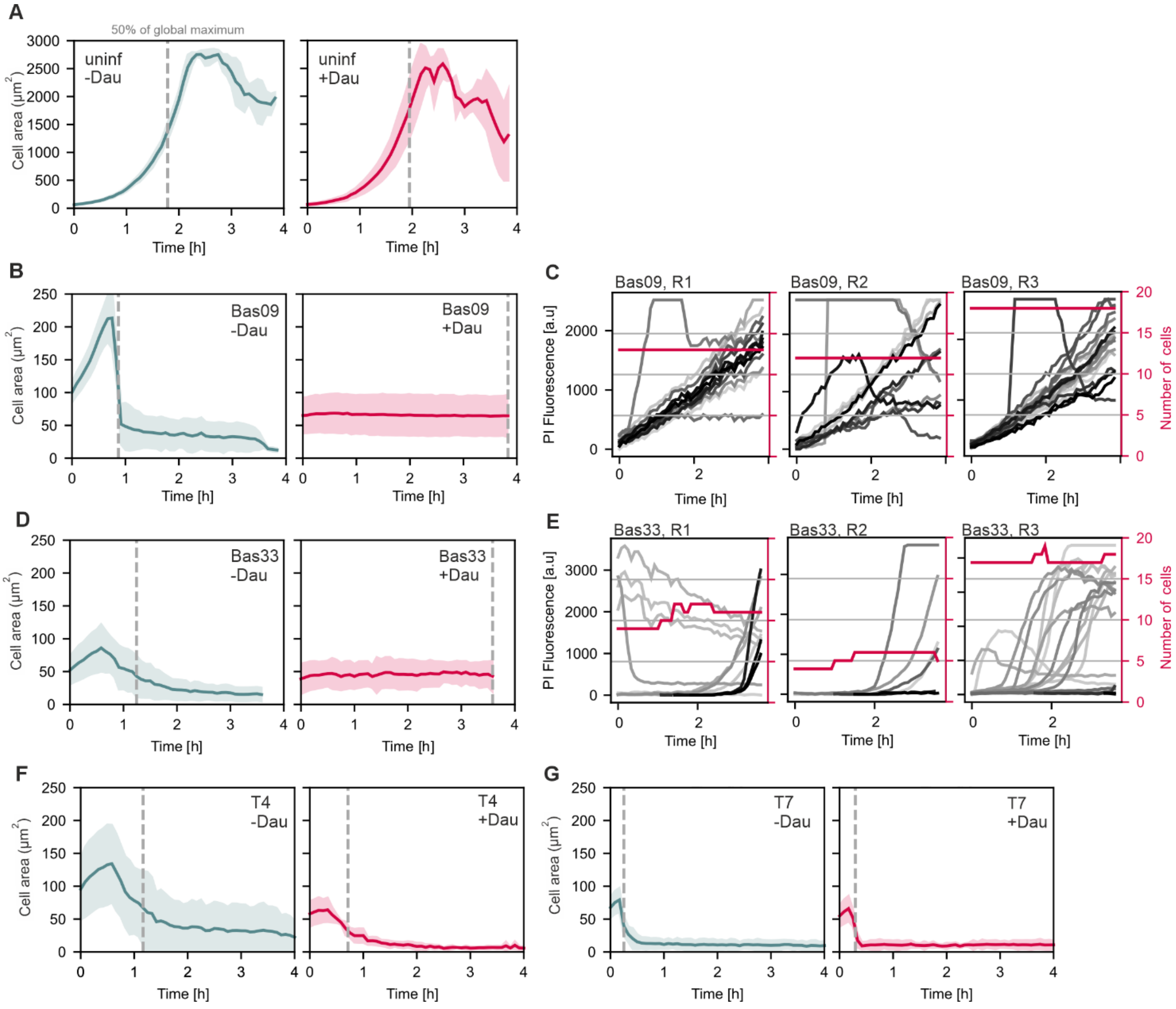


**Figure S2: Growth and fluorescence analysis on single-cell level during cultivation of *E. coli* K-12 MG1655 ΔRM in microfluidic chips with applied propidium iodide (PI) stain.** The dashed grey lines indicate the time points at which 50% of the global maximum cell area was reached. A) Mean of cell area during cultivation of *E. coli* K-12 MG1655 ΔRM in presence and absence of 2.5 µM daunorubicin. B) Cell area during infection of *E. coli* K-12 MG1655 ΔRM with Bas09 in presence and absence of 2.5 µM daunorubicin. C) PI fluorescence and number of cells upon infection of of *E. coli* K-12 MG1655 ΔRM with Bas09 in presence of 2.5 µM daunorubicin, shown for the single replicates. D) Cell area during infection of *E. coli* K-12 MG1655 ΔRM with Bas33 in presence and absence of 2.5 µM daunorubicin. E) PI fluorescence and number of cells upon infection of of *E. coli* K-12 MG1655 ΔRM with Bas33 in presence of 2.5 µM daunorubicin, shown for the single replicates. F) Cell area during infection of *E. coli* K-12 MG1655 ΔRM with T4 in presence and absence of 2.5 µM daunorubicin. G) Cell area during infection of *E. coli* K-12 MG1655 ΔRM with T7 in presence and absence of 2.5 µM daunorubicin. All analyses were performed as three independent biological replicates.

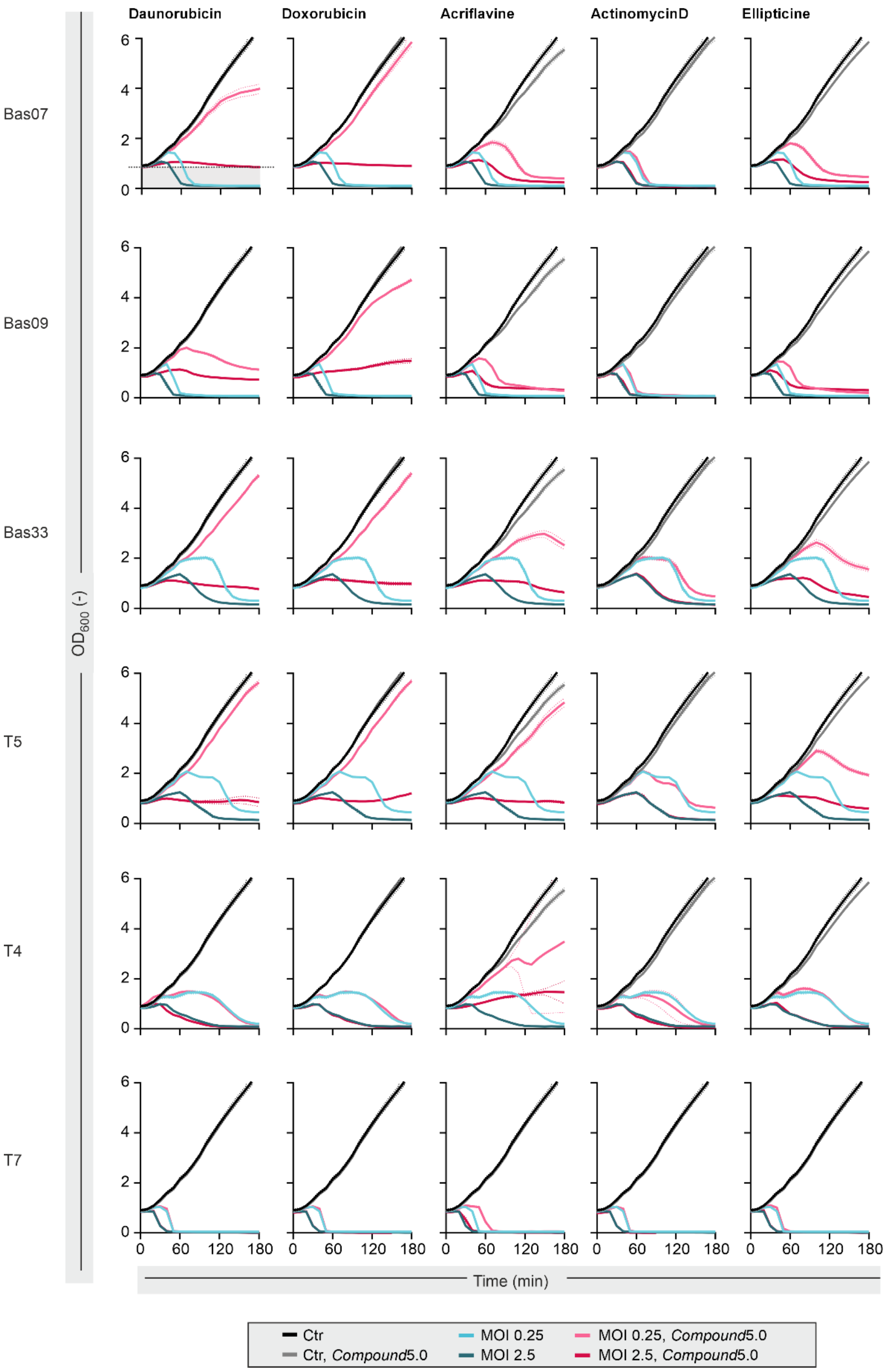


**Figure S3: Influence of different DNA-intercalating agents on phage infection dynamics.** Growth curves of *E. coli* K-12 MG1655 ΔRM upon infection with indicated phages in presence and absence of either 5 µM daunorubicin, doxorubicin, actinomycin D, ellipticine or 5 µg/ml acriflavine. All assays were performed as three independent biological replicates. For Bas07 and infection in presence of daunorubicin, principle of AUC calculation shown in Figure 2D is indicated by the dotted line showing the baseline (corresponds to $OD_{600}$ at t0) and the grey area showing the respective AUCs.

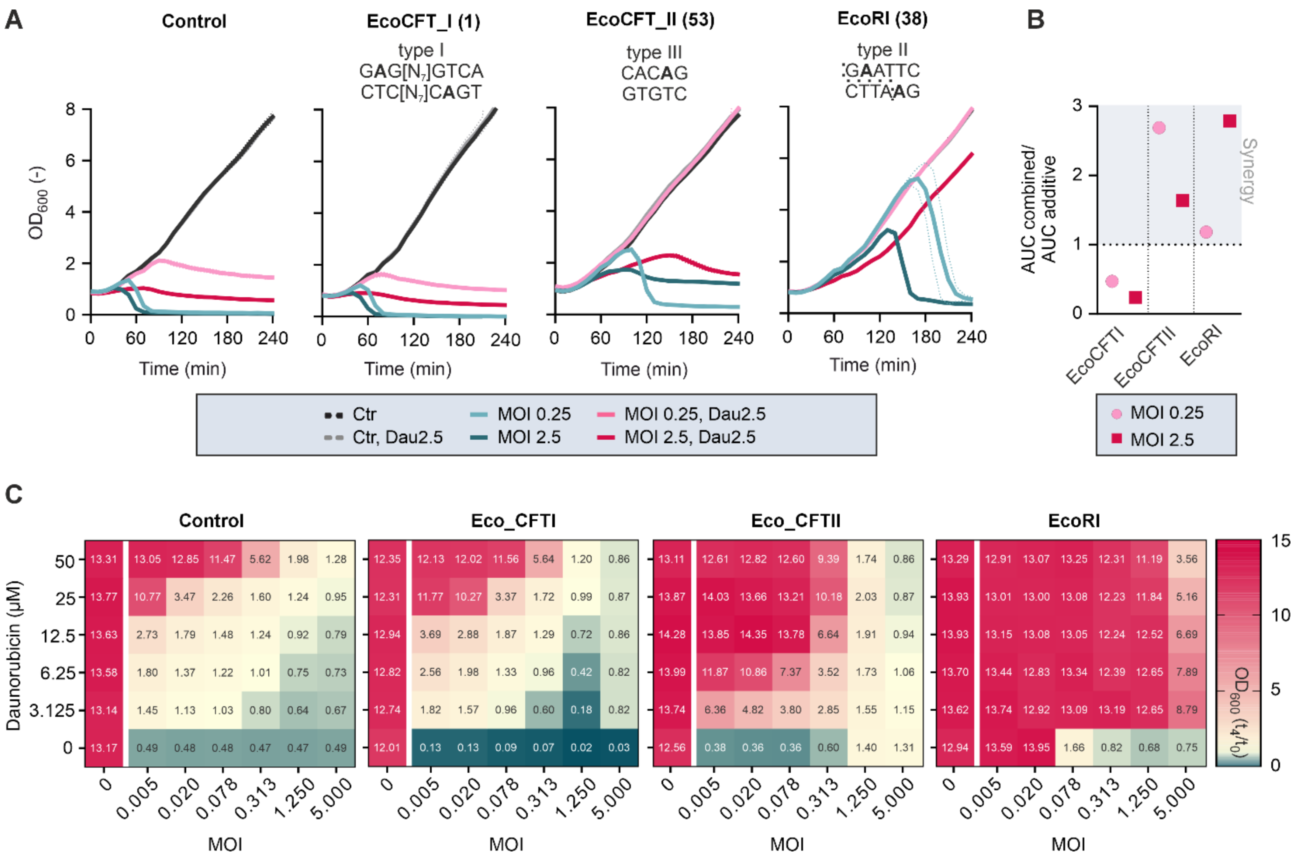


**Figure S1: DNA-targeting defense systems showed synergistic effects with daunorubicin towards Bas09 infection.** A) Liquid infection assays of the phage Bas09 infecting *E. coli* MG1655 ΔRM strains carrying different RM systems. Assays were performed in presence and absence of 2.5 µM daunorubicin at low and high MOIs of 0.25 and 2.5 (n =3). B) Calculation of synergy between daunorubicin and the respective RM systems based on the „Area under the curve (AUC)“ according to Wu et al. (2024) using $OD_{600}$ recorded at $t_0$ as baseline. C) Checkerboard-like assays combining different MOIs of the phage Bas09 and different daunorubicin concentrations in the respective *E. coli* strains with distinct RM antiphage defense backgrounds. The panel shows the FC in $OD_{600}$ after 4 h of cultivation and infection, with the calculated values indicated in the heatmaps.

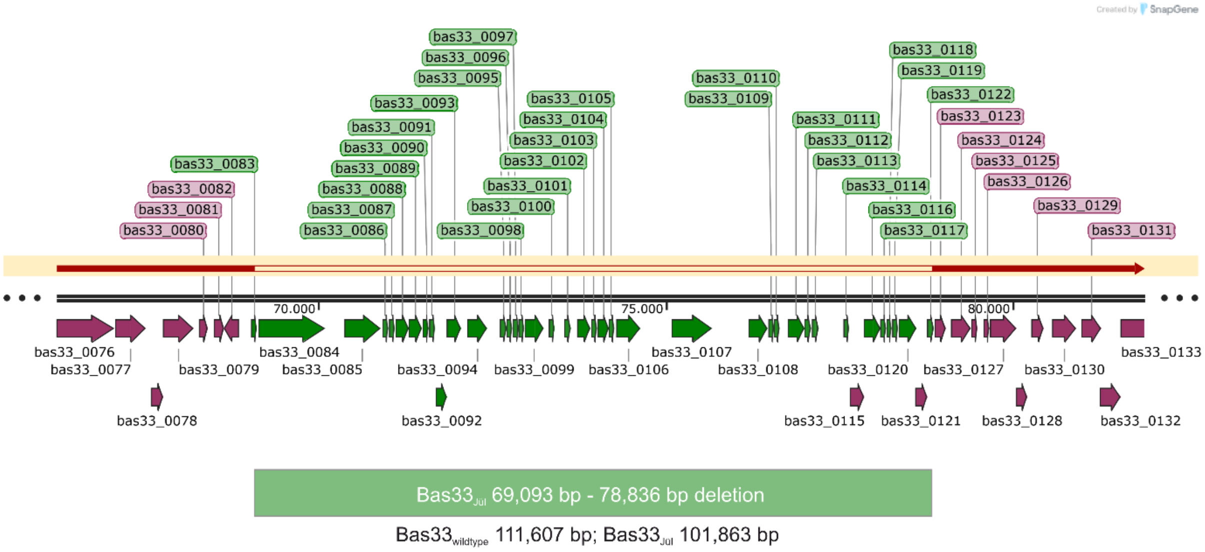


**Figure S5: Comparison of Bas33$_{wildtype}$ and Bas33$_{Jülich}$ genome.** The Bas33 variant used in all assays has a ~10 kbp deletion covering the basepairs 69,093 – 78,863. This region codes for several hypothetical proteins and tRNAs. A comparable deletion was already detected for T5$_{Mos}$ (Burman et al., 2024). However, the infection phenotype in the presence of daunorubicin was independently observed for several T5-like phages (Figure 1).

#### Supplementary Videos

**Video S1: Infection of *E. coli* K-12 MG1655 DRM with Bas33 in microfluidic chips using LB medium.** Microfluidic chambers were inoculated with *E. coli* cells and phages were added with the medium supply at a flow rate of 200 nl $min^{-1}$. Propidium iodide was added to visualize membrane permeabilization.

**Video S2: Infection of *E. coli* K-12 MG1655 DRM with Bas33 in microfluidic chips using LB medium with 2.5 µM daunorubicin.** Microfluidic chambers were inoculated with *E. coli* cells and phages were added with the medium supply at a flow rate of 200 nl $min^{-1}$. Propidium iodide was added to visualize membrane permeabilization.

**The videos are provided as separate files:**

- Video S1_Bas33 infection

- Video S2_Bas33 infection with 2.5 µM daunorubicin
